# A two-step approach to testing overall effect of gene-environment interaction for multiple phenotypes

**DOI:** 10.1101/2020.07.06.190256

**Authors:** Arunabha Majumdar, Kathryn S. Burch, Sriram Sankararaman, Bogdan Pasaniuc, W. James Gauderman, John S. Witte

## Abstract

While gene-environment (GxE) interactions contribute importantly to many different phenotypes, detecting such interactions requires well-powered studies and has proven difficult. To address this, we combine two approaches to improve GxE power: simultaneously evaluating multiple phenotypes and using a two-step analysis approach. Previous work shows that the power to identify a main genetic effect can be improved by simultaneously analyzing multiple related phenotypes. For a univariate phenotype, two-step methods produce higher power for detecting a GxE interaction compared to single step analysis. Therefore, we propose a two-step approach to test for an overall GxE effect for multiple phenotypes. Using simulations we demonstrate that, when more than one phenotype has GxE effect (i.e., GxE pleiotropy), our approach offers substantial gain in power (18% – 43%) to detect an aggregate-level GxE effect for a multivariate phenotype compared to an analogous two-step method to identify GxE effect for a univariate phenotype. We applied the proposed approach to simultaneously analyze three lipids, LDL, HDL and Triglyceride with the frequency of alcohol consumption as environmental factor in the UK Biobank. The method identified two independent genome-wide significant signals of an overall GxE effect on the vector of lipids.

## 1 Introduction

Gene-environment (GxE) interactions contribute significantly to the genetic architecture underlying complex phenotypes [1]. However, most GxE methods focus on testing a non-null effect of the interaction between one phenotype and one environmental factor at a time across genome-wide genetic variants [2,3], (e.g., approaches to jointly testing marginal and interaction effects [4], empirical Bayes shrinkage methods [5], two-step approaches [6–8], etc.). While these approaches can increase power to detect GxE interactions, adequate power remains a concern. One possible approach to further increase the power of detecting GxE interactions is by modeling multiple related phenotypes together. Previous work indicates that power to detect main genetic effects can be increased by modeling multiple correlated phenotypes; thus, one would expect similar gains to be available for assessing GxE interactions [9].

There exists substantial shared genetic basis among different phenotypes (i.e., pleiotropy). Genome-wide association studies (GWAS) have shown overlap in the main genetic effects across various complex phenotypes. While extensive work has investigated approaches for assessing pleiotropy in main genetic effects [10–16], little has been done with regard to assessing pleiotropy in GxE effects. For example, an interaction between physical activity and a genetic variant can influence the levels of three lipids, LDL, HDL and Triglycerides, simultaneously [17]. As another example, the pleiotropic genetic architecture of multiple smoking-related cancers (e.g., lung and head-neck) can be different among smokers and non-smokers [18, 19].

A recent study [20] proposed a mixed-model approach to quantify the heritability of a complex phenotype explained due to GxE interaction across multiple environmental factors for a single phenotype. Another study [21] proposed a subset-based multi-phenotype fixed-effects meta-analysis considering both marginal genetic effect and GxE effect across multiple phenotypes in the same model based on summary statistics of the corresponding effects. A simple strategy to test for an overall GxE effect across phenotypes is to perform a multivariate multiple linear regression considering both the multivariate main genetic effect and the interaction effect simultaneously in the model.

For a univariate phenotype, two-step methods can produce higher power for detecting GxE interactions compared to conventional approaches using a single analysis testing a GxE interaction. Two-step approaches filter out less important genetic variants in the first step and test the more promising variants for GxE interaction in the second step to reduce the multiple testing burden. Among various strategies in the first step, a common approach is to test the SNPs for a marginal genetic association with the phenotype under the assumption that a SNP having a GxE interaction effect on the phenotype should also have a marginal genetic effect on the phenotype. Similarly, a two-step procedure for multivariate phenotypes should produce higher power for detecting an aggregate-level GxE effect compared to a simple one-step multivariate regression of testing an overall GxE effect across phenotypes.

In this article, we extend the two-step procedure to multivariate quantitative phenotypes, and investigate its relative performance compared to the one-step multivariate regression for testing an overall GxE effect. Our motivation is two-fold: in the 1^*st*^ step, while filtering less important SNPs, simultaneously testing multiple related phenotypes should offer higher power for detecting SNPs having an overall marginal genetic effect (pleiotropy in main genetic effect); and in the 2^*nd*^ step, testing such selected promising SNPs for an aggregate-level GxE effect on the multiple phenotypes should produce higher power due to pleiotropy in GxE effect across the phenotypes.

To adjust for multiple testing in the one-step and two-step approaches, we considered three different procedures: Bonferroni correction, subset testing, and weighted hypothesis testing. We demonstrate by simulations that the multivariate two-step approach has a substantial power gain over the competing approaches. For real data application, we implement our approach to identify overall GxE effect of genome-wide SNPs and frequency of alcohol consumption on three lipids (LDL, HDL, Triglycerides) in the UK Biobank.

## 2 Methods

Let ***Y*** = (*Y*_1_, …, *Y_k_*)′ be multiple continuous phenotypes in a cohort, *G* denote genotypes at a SNP, and *E* an environmental factor. We consider multivariate linear regression (MLR) to model the main genetic effect of the SNP on *Y*.

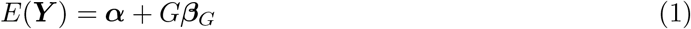

Here, 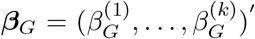 and ***α*** = (*α*^(1)^, …, *α*^(*k*)^)′, and the error component is assumed to follow a multivariate normal distribution with zero mean vector and covariance matrix Σ_1_. In the first step of the two-step procedure, we implement MLR to assess the overall main genetic effect of the SNP. In particular, we test 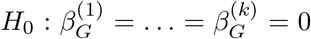 versus 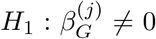, for at least one *j* = 1, …, *k*. We note that the power of identifying a SNP having a marginal genetic effect should improve by modeling multiple related phenotypes instead of a single phenotype. In the second step, we consider multivariate multiple linear regression (MMLR) to incorporate the multivariate main effects of the SNP (*G*) and the environmental factor (*E*), and the multivariate interaction effect due to GxE.

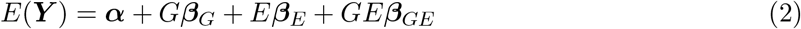

Here, 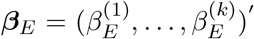 and 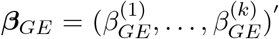, and the error component is assumed to follow multivariate normal with zero mean vector and a covariance matrix Σ_2_. We implement the type II MANOVA to test 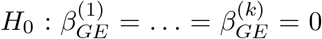 versus 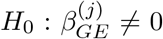, for at least one *j*. In the type II MANOVA test, the following two models are compared: the unrestricted full model, *E*(***Y***) = ***α*** + *G**β**_G_* + *E**β**_E_* + *GE**β**_GE_*, versus the restricted model, *E*(***Y***) = α + *G**β**_G_* + *E**β**_E_*. Here, the unrestricted model reduces to the restricted model under *H*_0_, when ***β**_GE_* = **0**. Thus, in the second step, we only test the null hypothesis that the vector of interaction effects ***β**_GE_* = **0**, leaving the vectors of main effects, ***β**_G_* and ***β**_E_*, unrestricted. The power of detecting a GxE interaction effect should be increased if the interaction is shared across *Y*_1_, …, *Y_k_*. We use the R package ‘car’ [22] to perform type II MANOVA.

In the two-step procedure, we combine the p-values obtained from 1^st^ and 2^nd^ steps to identify the SNPs that have a non-null overall GxE effect. We note that the linear model in equation (1) is nested under the linear model in equation (2). Hence, due to the general result in [6], the test statistic in the screening step to test ***β**_G_* = **0** (equation 1) and the test statistic testing ***β**_GE_* = **0** in the second step (equation 2) are independently distributed. This property is crucial to maintain the overall false positive rate of the combined two-step procedure at a desired level of significance. Below, we outline the subset testing (sst) approach and the weighted hypothesis testing (wht) approach to combine the two steps while adjusting for multiple testing to maintain the overall false positive rate.

### 2.1 Adjustment for multiple testing

Suppose we are considering *m* SNPs, and that we have two sets of p-values. One set is obtained from the first step testing for an overall main genetic effect across phenotypes (equation 1), 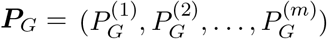 and the other set from the second step for the multivariate interaction effect (equation 2), 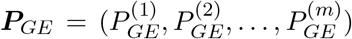. For the one-step multivariate GxE test, Bonferroni correction is applied to ***P**_GE_*. For the two-step approach, we consider the following two well-known procedures to combine the two steps [7, 23].

#### 2.1.1 Subset testing

For ***P**_G_*, we consider a p-value threshold *α*_1_ and filter out all SNPs for which 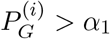, *i* = 1, …, *m*. In the second step, we only consider the SNPs selected in the first step (*P_G_* < *α*_1_), and apply a Bonferroni correction while testing for an overall GxE effect for these SNPs. Suppose, we consider a p-value threshold *α*_2_ in the second step. If *m*_1_ SNPs pass the first step, we compare *P_GE_* to 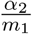 for each of the selected *m*_1_ SNPs to identify the SNPs having an overall GxE effect. A larger choice of *α*_1_ will increase the possibility of sending the SNPs with a true GxE effect to the second step, but at the expense of a higher multiple testing burden in the second step [7]. We considered a standard choice of the p-value thresholds: *α*_1_ = 0.05 and *α*_2_ = 0.05.

#### 2.1.2 Weighted hypothesis testing

Instead of completely dropping a set of less important SNPs in the second step, it has been argued that testing all SNPs in the second step while prioritizing them according to their relative ranking of importance obtained in the first step produces higher power to detect a GxE effect for a univariate phenotype [8,23,24]. Thus, we follow this approach and test all *m* SNPs using *P_GE_* in step 2 based on a significance level weighted using the order of the p-values in step 1 (*P_G_*). The weighting scheme uses an exponential weighting function, and allocates a larger fraction of the total significance level α to the most significant SNPs obtained in step 1 [8,23]. In particular, while performing step 2, the *k*_1_ most significant SNPs in the first bin in step 1 (lowest *P_G_*) are tested at a significance level 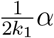, the next *k*_2_ (= 2*k*_1_) most significant SNPs in the second bin in step 1 are tested at 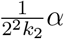, the next *k*_3_ (= 2*k*_2_) at 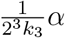, and so on [23]. For example, when *k*_1_ = 5 and *α* = 0.05, the top 5 SNPs from step 1 are tested at a significance level 0.005 in step 2, the next 10 at 0.00125, etc. This weighting scheme guarantees that the overall false positive rate for the entire procedure does not exceed *α*. Under this weighting scheme, the top SNPs from step 1 are tested at a more liberal significance threshold than the standard Bonferroni-corrected level required in a standard one-step exhaustive scan of all *m* SNPs. However, for the SNPs not in the top bins in step 1, weighted testing can have a more stringent threshold than Bonferroni correction. We used a standard choice of *k*_1_ = 5 and *α* = 0.05 [8,23].

### 2.2 GxE tests for univariate phenotype

To test for a GxE interaction for a univariate phenotype, we consider the following existing methods [6–8]. Let *Y* denote a single continuous phenotype. In the one-step approach, we consider *E*(*Y*) = *α* + *β_G_* × *G* + *β_E_* × *E* + *β_GE_* × *GE*, and test for *H*_0_: *β_GE_* = 0 versus *H*_1_: *β_GE_* ≠ 0. In the two-step approach, we combine the step 1 model: *E*(*Y*) = *α* + *β_G_* × *G*, with step 2 model: *E*(*Y*) = *α* + *β_G_* × *G* + *β_E_* × *E* + *β_GE_* × *GE*. Here we consider the same multiple testing strategies as considered above for a multivariate phenotype.

## 3 Simulation study

### 3.1 Framework

We describe the simulation design for two phenotypes mainly for convenience in presenting mathematical expressions. This can be extended for a larger number of phenotypes in a straightforward manner. Let *Y*_1_ and *Y*_2_ denote two phenotypes, *G* denote the genotypes at a SNP, and *E* an environmental factor. We consider the following bivariate multiple linear regression to model the phenotypes.

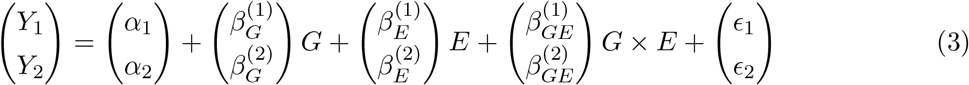

We consider each of *Y*_1_, *Y*_2_, *G* and *E* to be mean-centered. We assume a bivariate normal distribution for (*ϵ*_1_, *ϵ*_2_)′. Under a fixed effects model, 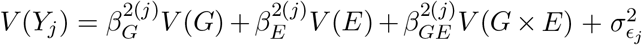, *j* = 1,2. Under the assumption that *G* and *E* are independent in the population, we obtain that *V*(*G* × *E*) = *V*(*G*)*V*(*E*), since *E*(*G*) = *E*(*E*) = 0. Thus, 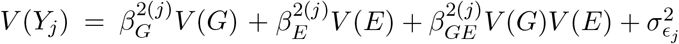, for *j* = 1, 2.

Let us denote 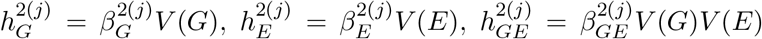, and the total variance of *j^th^* phenotype as 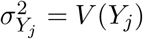. Hence, 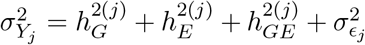. Without loss of generality, we assume that 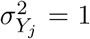; so, 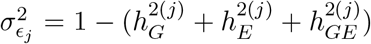; *j* = 1,2. Next, we derive the following: 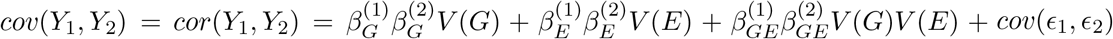, where *cov*(*ϵ*_1_, *ϵ*_2_) is the covariance between the noise terms in the two phenotypes. Thus, we can fix the correlation between the phenotypes (*cor*(*Y*_1_, *Y*_2_)) and other simulation parameters which in turn determine the value of *cov*(*ϵ*_1_, *ϵ*_2_) to be used in the simulations.

### 3.2 Choice of parameters

In our simulation study, we consider three phenotypes for 20,000 individuals, and choose the pairwise phenotypic correlations randomly in the range 20% – 30%. We consider 100, 000 null SNPs which have no marginal genetic association with any phenotype, and no GxE interaction on any phenotype. We simulate the minor allele frequency at a SNP from Uniform(0.05, 0.45), and simulate the genotypes under Hardy-Weinberg equilibrium (HWE). We consider a separate set of 100 non-null SNPs (denoted by *m_G_*) each of which has a marginal genetic effect on at least one phenotype (equation 1). Among these *m_G_* non-null SNPs, *m_GE_* SNPs have a GxE effect on at least one phenotype (equation 2), and we vary *m_GE_* = 10, 20, 30, 40. So, a subset of the risk SNPs having a marginal genetic effect are assumed to have a GxE effect (*m_GE_* out of *m_G_* = 100). We further assume that, if a SNP has a GxE effect on a phenotype, the same phenotype also has a marginal genetic effect due to the SNP. We consider three different scenarios. In the 1^st^ scenario, each non-null SNP has a marginal genetic effect on the first phenotype but not the other two phenotypes; and if the SNP (one of *m_GE_* SNPs) has a GxE effect, it has the interaction effect only on the first phenotype. Similarly in the 2^nd^ scenario, the first two phenotypes (but not the last phenotype) have a marginal genetic effect due to each non-null SNP, and each of *m_GE_* SNPs has a GxE effect on the first two phenotypes but not the last one. And in the 3^rd^ scenario, all the three phenotypes have a marginal genetic effect from each non-null SNP, and each of *m_GE_* SNPs has a GxE effect on every phenotype.

Under each scenario, we compare the performance of various tests of GxE interaction for each univariate phenotype and the multivariate phenotype. We apply three different procedures for multiple testing adjustment to control the family-wise error rate (FWER) as outlined above: Bonferroni correction for the one-step methods (abbreviated as bonf), subset testing (sst), and weighted hypotheses testing (wht). For each method, we estimate the type I error rate and power based on 200 simulated datasets. Under a given simulation scenario, for each simulated dataset comprising 10^5^ null SNPs and 100 risk SNPs, we compute the proportion of null SNPs at which the method of choice to test GxE (null hypothesis of no overall/univariate GxE effect) wrongly identified a genome-wide significant signal of interaction. We estimate the type I error rate as the mean of this proportion across 200 simulated datasets. We estimate the power using a similar procedure based on the risk SNPs only. For ease of presentation, when plotting the power obtained by the tests of GxE interaction for univariate phenotypes, we plot the maximum power obtained across three univariate phenotypes for each multiple testing adjustment procedure. This allows us to explore whether we obtain higher power by testing GxE interaction for a multivariate phenotype compared to each univariate phenotype.

### 3.3 Results

We present the estimated overall type I error rate obtained from GxE tests for a multivariate phenotype in Table 1. While the type I error rate appears to be controlled overall at the desired level of significance 0.05 with per-SNP level 5 × 10^−7^ (for 10^5^ null SNPs), we observe marginal inflation in some cases; this is mainly due to using 200 iterations of simulation (for computational feasibility) to estimate the FWER under a given simulation scenario. We note that the weighted hypothesis testing (our preferred choice of multiple testing strategy for a *2-step* procedure) well controls the type I error rate for the majority of the simulation choices (Table 1). We present the estimated type I error rate of GxE tests for univariate phenotypes in Table 2, and find that the false positive rate is controlled overall with marginal inflation in some cases. Here, the *2-step* procedure based on weighted hypothesis testing appears to better control the overall type I error rate (Table 2).

**Table 1:** Simulation results: estimated overall type I error rate obtained by different tests of overall GxE effect for a multivariate phenotype using various strategies of multiple testing adjustment. Here, #asso-pheno denotes the number of phenotypes that have a marginal genetic effect or an interaction effect due to risk SNPs; *m_GE_* denotes the number of SNPs out of 100 (*m_G_*) risk SNPs that have a GxE interaction effect. We present the overall type I error rate (FWER) at 0.05 level of significance with the desired level 5 × 10^−7^ per SNP, since we considered 10^5^ null SNPs which have no marginal G or GxE effect. The type I error rate is estimated based on 200 simulated datasets.

| #asso-pheno | $m_{GE}$ | *1-step* Bonf. correction | *2-step* subset testing | *2-step* weighted multiple testing |
| --- | --- | --- | --- | --- |
| 1 | 10 | 0.07 | 0.04 | 0.05 |
| 1 | 20 | 0.095 | 0.05 | 0.08 |
| 1 | 30 | 0.05 | 0.02 | 0.04 |
| 1 | 40 | 0.07 | 0.03 | 0.05 |
| 2 | 10 | 0.06 | 0.07 | 0.06 |
| 2 | 20 | 0.06 | 0.05 | 0.05 |
| 2 | 30 | 0.07 | 0.06 | 0.04 |
| 2 | 40 | 0.03 | 0.03 | 0.03 |
| 3 | 10 | 0.04 | 0.06 | 0.04 |
| 3 | 20 | 0.05 | 0.06 | 0.04 |
| 3 | 30 | 0.06 | 0.07 | 0.03 |
| 3 | 40 | 0.05 | 0.08 | 0.04 |

**Table 2:**
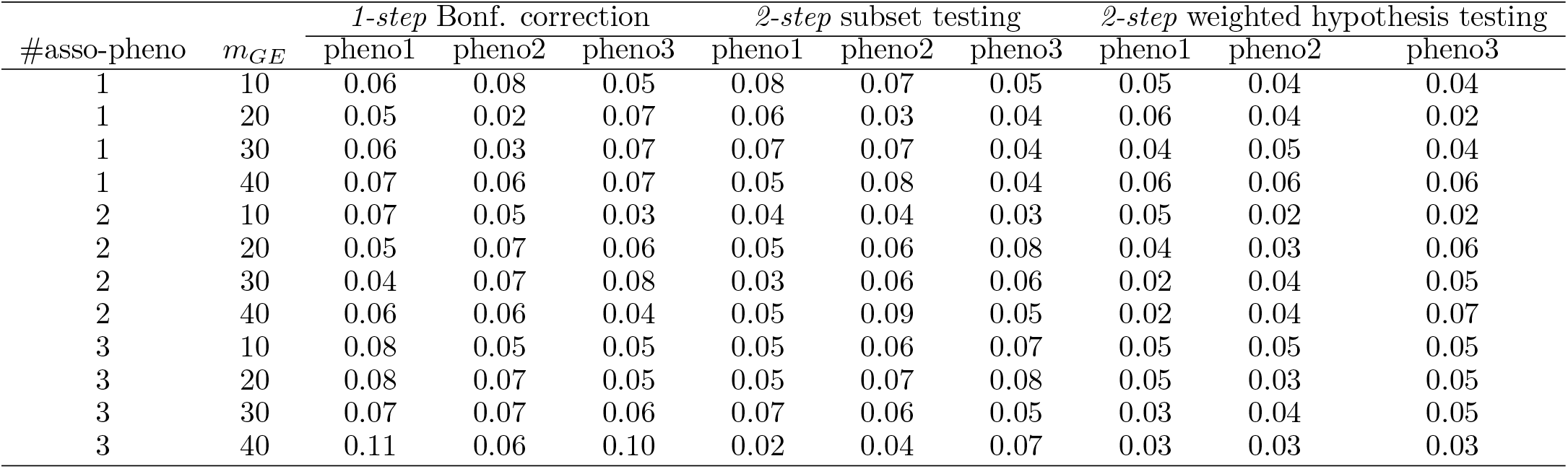
Simulation results: estimated overall type I error rate obtained by different tests of GxE effect for univariate phenotype using various strategies of multiple testing adjustment. Here, #asso-pheno denotes the number of phenotypes that have a marginal genetic effect or an interaction effect due to risk SNPs; *m_GE_* denotes the number of SNPs out of 100 (*m_G_*) risk SNPs that have a GxE interaction effect. We present the overall type I error rate (FWER) at 0.05 level of significance with the desired level 5 × 10^−7^ per SNP, since we considered 10^5^ null SNPs which have no marginal G or GxE effect. Three univariate phenotypes are abbreviated as pheno1, pheno2 and pheno3, respectively. The type I error rate is estimated based on 200 simulated datasets.

We present the estimated power of GxE tests for multivariate and univariate phenotypes in Figure 1, 2 and 3. First, we focus on the multivariate phenotype, and compare the power of *1-step* and *2-step* approaches to detect an overall effect of GxE interaction on multiple phenotypes. We find that both of the *2-step* procedures (subset testing and weighted hypothesis testing) produce higher power than the *1-step* approach (Bonferroni correction). We also observe that the weighted hypothesis testing (wht) performs better than the subset testing (sst). We therefore focus on comparing the weighted hypothesis testing procedure with the Bonferroni correction to contrast the power of *2-step* and *1-step* approaches. In 3^rd^ simulation scenario, when all three phenotypes have a GxE effect from each of *n_GE_* SNPs, the *2-step* approach implemented by weighted hypothesis testing (abbreviated as *2-step*-wht) produces 31%–34% higher power than the Bonferroni correction implementing the *1-step* approach (*1-step*-bonf) (Figure 3). In 2^nd^ scenario, when two phenotypes have a GxE effect, *2-step*-wht produces 23% – 24% power increase than *1-step*-bonf (Figure 2). In the absence of pleiotropy in GxE effect, i.e., when one phenotype has a GxE effect, *2-step*-wht yields 7% – 8% power gain than *1-step*-bonf (Figure 1). Hence, overall our *2-step* approach offers substantial power gain compared to the *1-step* approach when testing for an overall GxE effect on a multivariate phenotype. We also find that *2-step*-wht performs consistently better than *2-step*-sst (subset testing) and produces a power gain of 16% – 18% in the third simulation scenario (Figure 3), 14% – 15% in the second scenario (Figure 2), and 6% in the first scenario (Figure 1).

**Figure 1:**
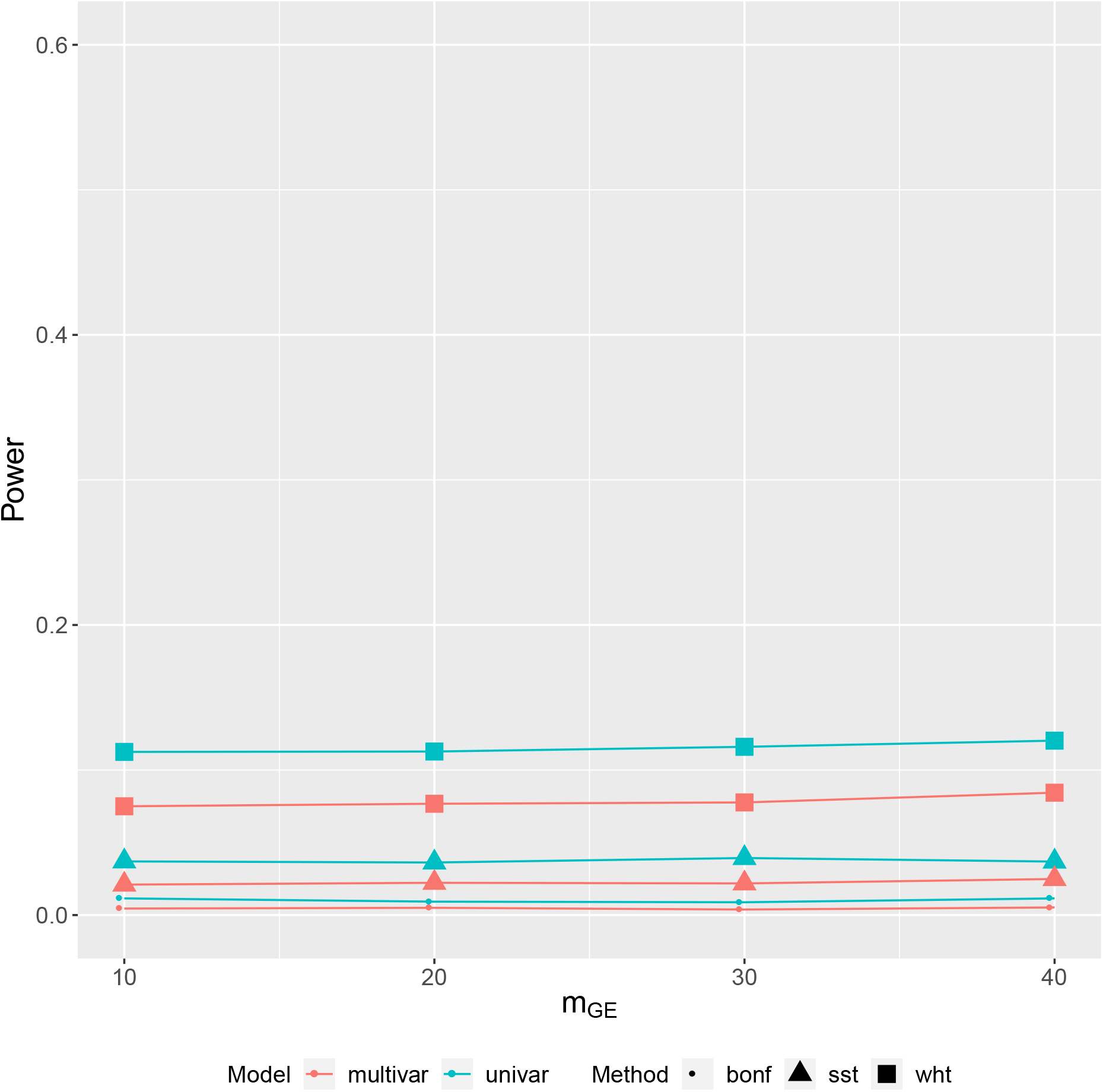
Simulation results: estimated power obtained by different tests of overall GxE effect for multivariate phenotype (multivar), and tests of GxE effect for univariate phenotype (univar) using various strategies of multiple testing adjustment: *1-step* Bonferroni correction (bonf), *2-step* subset testing (sst), and *2-step* weighted hypothesis testing (wht). Here, 1^st^ phenotype (but not 2^nd^ and 3^rd^) has a marginal genetic effect or an interaction effect due to risk SNPs. We denote the number of SNPs out of 100 risk SNPs which have a GxE effect as *m_GE_*. The power is estimated based on 200 simulated datasets.

**Figure 2:**
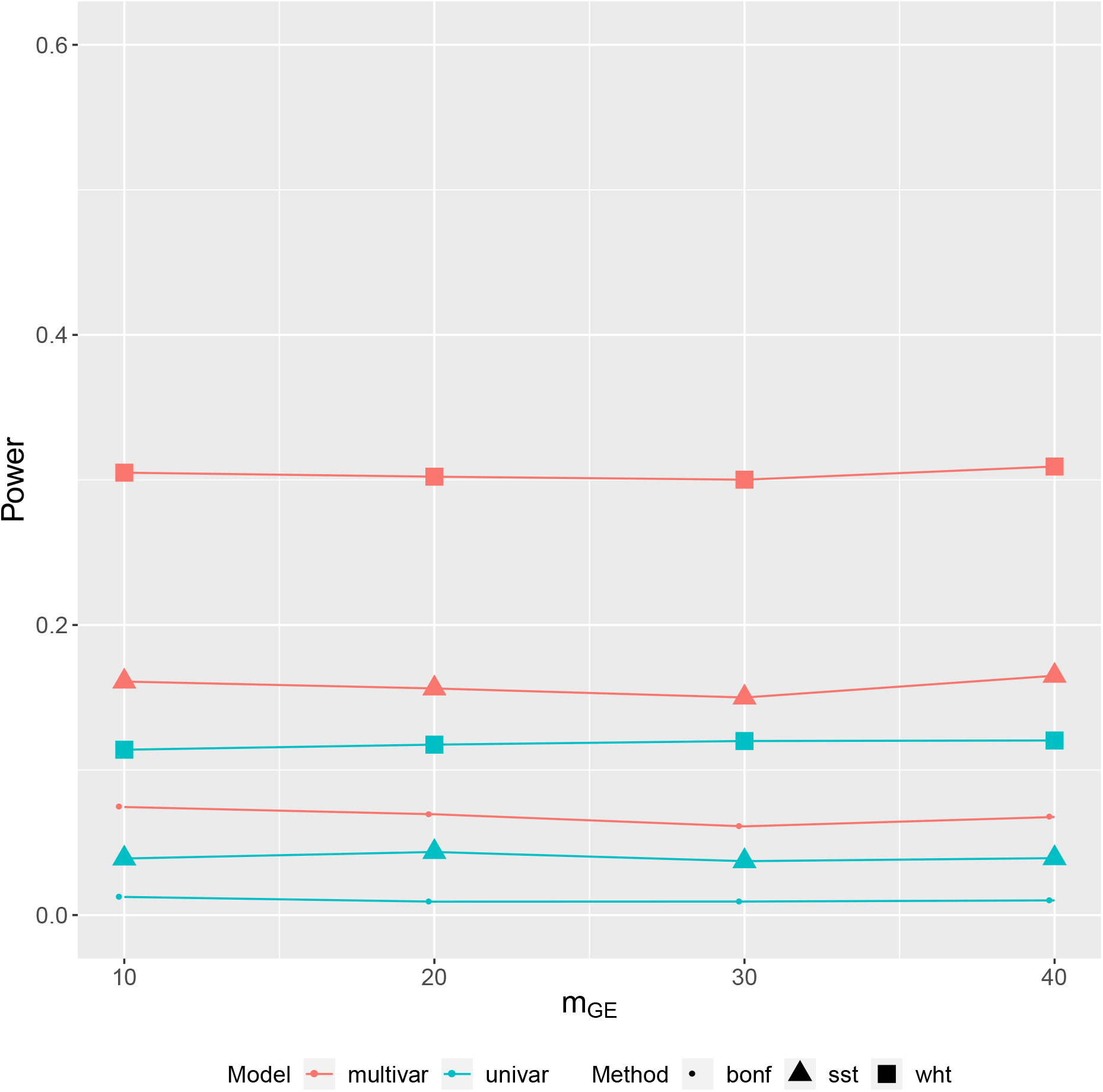
Simulation results: estimated power obtained by different tests of overall GxE effect for multivariate phenotype (multivar), and tests of GxE effect for univariate phenotype (univar) using various strategies of multiple testing adjustment: *1-step* Bonferroni correction (bonf), *2-step* subset testing (sst), and *2-step* weighted hypothesis testing (wht). Here, first two phenotypes (but not 3^rd^) have a marginal genetic effect or an interaction effect due to risk SNPs. We denote the number of SNPs out of 100 risk SNPs which have a GxE effect as *m_GE_*. The power is estimated based on 200 simulated datasets.

**Figure 3:**
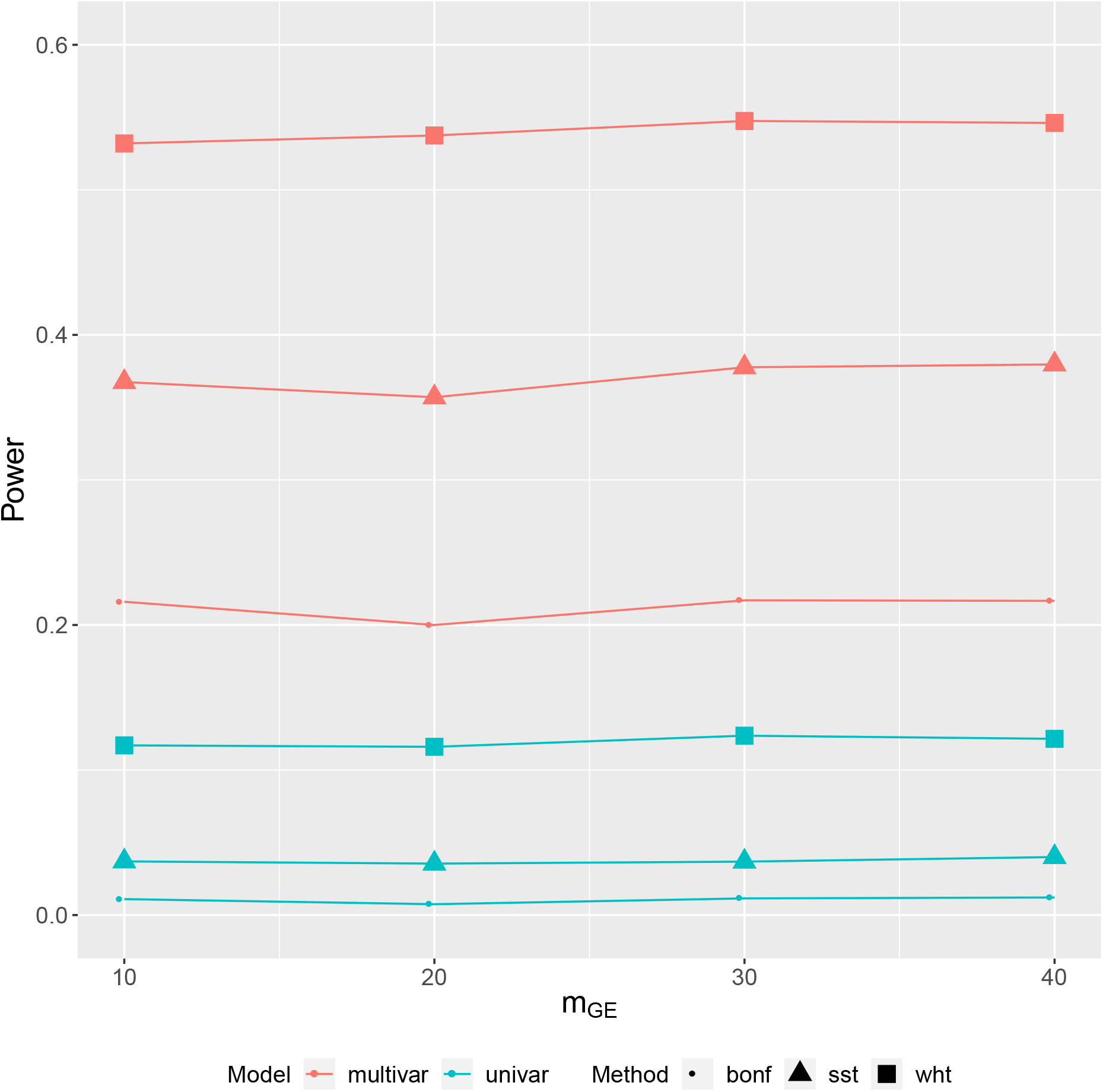
Simulation results: estimated power obtained by different tests of overall GxE effect for multivariate phenotype (multivar), and tests of GxE effect for univariate phenotype (univar) using various strategies of multiple testing adjustment: *1-step* Bonferroni correction (bonf), *2-step* subset testing (sst), and *2-step* weighted hypothesis testing (wht). Here, all three phenotypes have a marginal genetic effect or an interaction effect due to risk SNPs. We denote the number of SNPs out of 100 risk SNPs which have a GxE effect as *m_GE_*. The power is estimated based on 200 simulated datasets.

For GxE tests with univariate phenotypes, the *2-step* approaches (wht and sst) performed better than the *1-step* approach (bonferroni correction) which is consistent with findings from previous studies [7, 8]. Between the two strategies of *2-step* approaches for univariate phenotype, wht produces higher power than sst (Figure 1, 2 and 3).

Next, we contrast the performance of *2-step*-wht for multivariate phenotypes (multivar_wht) with that of *2-step*-wht for univariate phenotypes (univar_wht). For ease of comparison, we contrast the power obtained by multivar_wht with the maximum power obtained across the three univariate phenotypes each obtained by univar_wht. In the presence of pleiotropy in the GxE effect, multivar_wht produces 41% – 43% higher power than univar_wht under the third simulation scenario (Figure 3), and 18% – 20% power gain under the second scenario (Figure 2). However, in the absence of pleiotropy in GxE effect under the first simulation scenario, univar_wht offers marginal power gain (3% – 4%) over multivar_wht (Figure 1). Taken together, the multivariate approach produces substantially higher power than the univariate approach in the presence of pleiotropy in GxE effect, and loses marginal power in its absence.

## 4 Real data application

We considered three lipids LDL, HDL, and Triglycerides in the UK Biobank as the multivariate phenotype, and the frequency of alcohol consumption as the environmental factor. We removed individuals with missing values of phenotypes or relevant covariates from the sample, leaving 253,653 White-British unrelated individuals. First, we apply the inverse rank normal transformation on each lipid separately; then adjust each transformed phenotype for age, sex, and 20 principal components (PCs) of genetic ancestry by linear regression. We consider the adjusted residual from each linear regression as the final phenotype vectors *Y_j_, j* = 1,2,3. We tested 459, 792 genotyped SNPs in the UKB one at a time. We applied the different procedures presented above for testing the GxE interaction. We provide the results from multivariate tests of GxE interaction in Table 3.

**Table 3:** Real data results: genome-wide significant signals of aggregate-level GxE interaction effect on the vector of three lipids (LDL, HDL, Triglycerides) obtained by multivariate tests for overall GxE effect. The frequency of alcohol consumption is considered as the environmental factor. CHR denotes chromosome, and BP denotes base pair position. P.g denotes the p-value of testing the multivariate marginal genetic association between the SNP and the lipids; P.ge (MMLR) denotes the p-value of testing overall GxE effect on the lipids using multivariate multiple linear regression (MMLR) prior to adjustment for multiple testing; P.ge.wht denotes the p-value of testing the overall GxE effect using the *2-step* approach based on weighted hypothesis testing. The lead SNPs, i.e., the independent SNPs having the strongest genome-wide significant signal of aggregate-level GxE effect, are colored blue. The lead SNPs are obtained based on *r*^2^ threshold 0.2.

| *2-step* weighted hypothesis testing |  |  |  |  |  |
| --- | --- | --- | --- | --- | --- |
| SNP | CHR | BP | P.g | P.ge (MMLR) | P.ge.wht |
| rs6984305 | 8 | 9178268 | 6.09E-111 | 6.12E-08 | 0.0006 |
| rs11782386 | 8 | 9201787 | 1.35E-64 | 3.89E-06 | 0.04 |
| rs11779870 | 8 | 9211723 | 5.34E-68 | 4.80E-06 | 0.05 |
| rs964184 | 11 | 116648917 | < E-300 | 6.86E-05 | 0.01 |
| *2-step* subset testing |  |  |  |  |  |
|  |  |  | P.g | P.ge (MMLR) |  |
| rs6984305 | 8 | 9178268 | 6.09E-111 | 6.12E-08 |  |
| *1-step* Bonferroni correction |  |  |  |  |  |
|  |  |  | P.g | P.ge (MMLR) |  |
| rs77400814 | 2 | 24073810 | 0.41 | 1.26E-08 |  |
| rs76614804 | 2 | 24626991 | 0.70 | 6.63E-10 |  |
| rs6984305 | 8 | 9178268 | 6.09E-111 | 6.12E-08 |  |

For four SNPs on chromosome 8 and 11, the *2-step* weighted hypothesis testing (*2-step*-wht) identified a genome-wide significant overall effect of GxE interaction on the lipids, whereas the *1-step* Bonferroni correction (*1-step*-bonf) identified an overall interaction effect for three SNPs on chromosome 2 and 8 (Table 3). However, the *2-step* subset testing (*2-step*-sst) identified an overall GxE effect for only one SNP on chromosome 8 (rs6984305 which was also identified by *2-step*-wht and 1-step-bonf). All the SNPs on chromosome 8 detected by *2-step*-wht are in linkage disequilibrium (LD). We identified the lead SNP (colored blue) based on *r*^2^ threshold of 0.2 (Table 3). Similarly, *1-step*-bonf detected a pair of SNPs on chromosome 2 which are in LD. We note that *2-step*-wht detected a signal on chromosome 11 which was missed by 1-step-bonf. However, *2-step*-wht missed the signal on chromosome 2 identified by *1-step*-bonf (Table 3).

For simplicity, results from the univariate analysis based on *2-step* weighted hypothesis testing (*2-step*-wht) procedure are presented in Table 4. In simulations, *2-step*-wht produced highest power for univariate case. For HDL, it identified genome-wide significant signal of GxE effect, but none for LDL and triglycerides. Even though it detected six SNPs on chromosome 8 for HDL, all of them are in strong LD resulting in the same lead SNP rs6984305, which was also identified by the multivariate tests (Table 3). We note that the multivariate *2-step*-wht identified the GxE signal on chromosome 11 (rs964184) which was missed by the analogous univariate *2-step*-wht.

**Table 4:** Real data results: genome-wide significant signals of univariate GxE effect on HDL obtained by *2-step* weighted hypothesis testing. The frequency of alcohol consumption is considered as the environmental factor. P.g denotes the p-value of testing univariate marginal genetic association between the SNP and HDL; P.ge (UMLR) denotes the p-value of testing univariate GxE effect on HDL using univariate multiple linear regression (UMLR) prior to adjustment for multiple testing; P.ge.wht denotes the p-value of testing univariate GxE effect using *2-step* weighted hypothesis testing. The lead SNPs, i.e., the independent SNPs having the strongest genome-wide significant signal of univariate GxE effect, are colored blue. The lead SNPs are obtained based on *r*^2^ threshold 0.2.

| SNP | CHR | BP | P.g | P.ge (UMLR) | P.ge.wht |
| --- | --- | --- | --- | --- | --- |
| rs6984305 | 8 | 9178268 | 1.61E-78 | 1.40E-08 | 3.59E-05 |
| rs9987289 | 8 | 9183358 | 1.87E-98 | 4.12E-06 | 0.003 |
| rs6601299 | 8 | 9184691 | 5.29E-79 | 3.28E-06 | 0.008 |
| rs2126259 | 8 | 9185146 | 4.52E-81 | 2.73E-06 | 0.007 |
| rs11782386 | 8 | 9201787 | 1.40E-48 | 2.86E-06 | 0.007 |
| rs11779870 | 8 | 9211723 | 5.27E-51 | 2.68E-06 | 0.007 |

At rs964184 on chromosome 11, the univariate GxE test (multiple linear regression) p-value for three lipids were 0.004, 0.0007 and 0.009, none of which is genome-wide significant. Even though this seems to be a moderate evidence of pleiotropy in GxE effect, the *1-step* multivariate test (MMLR) p-value for an overall GxE effect across lipids was 6.9 × 10^−5^ which is also not genome-wide significant. However, since this SNP has a strong evidence of pleiotropy in marginal main genetic effect with univariate p-values across lipids as 5.8 × 10^−52^, 2.6 × 10^−176^, P < 10^−300^, and also a p-value < 10^−300^ for the multivariate main genetic association, the multivariate *2-step*-wht approach prioritized this SNP in the second step while testing GxE, and identified a genome-wide significant overall GxE effect for this SNP.

At rs6984305 on chromosome 8 which is mapped to a nearby gene AC022784.1, a previous study [25] identified an effect of GxE interaction on HDL with sleep duration as the environmental factor. In NHGRI-EBI GWAS catalog, this SNP is also reported to be marginally associated with liver enzyme levels and serum alkaline phosphatase levels. At rs964184 on chromosome 11, previous studies found GxE interaction effect on HDL with sleep duration as the environmental factor [25], on triglycerides with physical activity as the environmental factor [17]. It was also reported to be marginally associated with many phenotypes including lipid levels, cardiovascular disease, red blood cell phenotypes, etc.

## 5 Discussion

We have proposed a two-step approach to test for an aggregate-level gene-environment interaction across multiple related phenotypes. Using simulations, we demonstrate that our method produces substantially higher power than the Bonferroni-corrected one-step test of overall effect of GxE interaction on a multivariate phenotype. While our proposed approach also provides substantially higher power than competing univariate approaches in the presence of pleiotropic GxE effect, in the absence of pleiotropy, the method only loses marginal power compared to the analogous two-step univariate approach.

We demonstrate our *2-step* approach by applying it to a vector of three lipid phenotypes in the UK Biobank with the frequency of alcohol consumption as the environmental factor. Our method identified a pair of independent genome-wide significant signals of overall effect of GxE interaction on the three lipids. Previous studies reported these SNPs to have GxE effect on HDL with sleep duration as the environmental factor, and on triglycerides with physical activity as the environmental factor.

There are some limitations to our approach and potential for future improvement. First, as commonly practiced in standard GWAS, the multiple testing procedures implemented in our approach to identify the genome-wide significant signals of GxE effect do not explicitly account for LD among the SNPs. Hence, the procedures are expected to be conservative in nature, limiting the power of the tests. An interesting future direction of research will be to adjust the multiple testing strategies for LD, in particular, the weighted hypothesis testing and subset testing procedures. Second, we have only explored the performance of our proposed approach for quantitative phenotypes. In future work, we plan to extend the approach for multiple related case–control phenotypes under a logistic regression framework. Third, we have considered one environmental factor in the model. However, considering multiple relevant environmental factors at the same time as multiple phenotypes should further improve the power of detecting an overall GxE effect.

In summary, our proposed *2-step* approach is a powerful method to detect an overall effect of GxE interaction on a multivariate phenotype. The approach is theoretically sound and computationally efficient. It can be implemented using the existing R software package ‘car’.

## 6 Acknowledgement

This research was conducted using the UK Biobank Resource under applications 24129 and 33297. We thank the participants of UK Biobank for making this work possible.

